# Nrf2 mediated ER-phagy protects against oxidative damage in intervertebral disc degeneration

**DOI:** 10.1101/2021.11.21.469451

**Authors:** Zhen Lin, Libin Ni, Cheng Teng, Zhao Zhang, Xinlei Lu, Chenglong Xie, Long Wu, Yifei Zhou, Naifeng Tian, Yaosen Wu, Liaojun Sun, Zongyou Pan, Xiangyang Wang, Zhongke Lin, Xiaolei Zhang

## Abstract

Intervertebral disc degeneration (IDD) increases the risk of low back pain (LBP). Oxidative stress may induce cellular damage and contribute to various diseases including IDD. Endoplasmic reticulum autophagy (ER-phagy) is a specific type of autophagy, its role in oxidative stress induced damage as well as in IDD is unknown. This study explores the role of ER-phagy in oxidative damage in intervertebral disc nucleus pulposus cells (NPCs), as well as the Nrf2/FAM134B axis in ER-phagy regulation and IDD therapy. We found ER-phagy was decreased in NPCs during oxidative stress; while FAM134B may promote ER-phagy and alleviate oxidative stress induced ER-stress and apoptosis. In addition, the nuclear transcription factor Nrf2 may promote the expression of FAM134B as well as ER-phagy, and suppress ER-stress and apoptosis in NPCs. Furthermore, overexpression of FAM134B and Nrf2 could effectively attenuate the progression of IDD in rats *in vivo*. These results suggest Nrf2/FAM134B mediated ER-phagy may combat oxidative damage in cells; meanwhile, ER-phagy as well as Nrf2 could be potential therapeutic targets for IDD.

**Graphical abstract:** 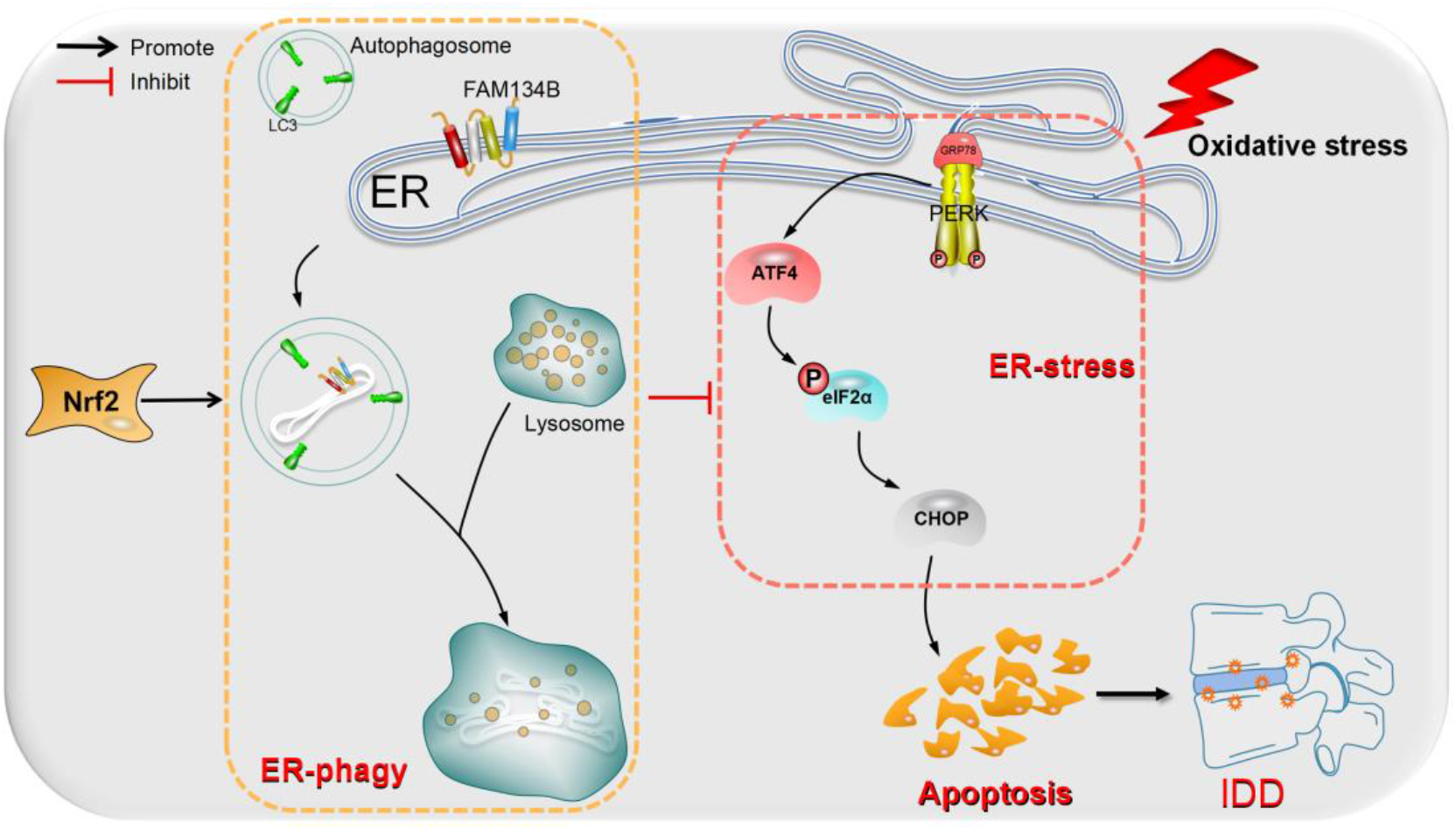

## 1. Introduction

Low back pain (LBP) affects nearly 80% of the population worldwide. It not only reduces the quality of daily life, but also lead to disability in severe cases, and cause heavy economic burden on a global scale(Deyo, Loeser, & Bigos, 1990). Among the many influencing factors leading to its occurrence, intervertebral disc degeneration (IDD) has been proven to be one of the most important causes(Vergroesen et al., 2015).

Although the pathophysiological mechanism of the occurrence and development of IDD is complicated; recently, increasing evidence has verified the presence of oxidative stress and increased concentrations of oxidation products in IDD(Feng et al., 2017; Scharf et al., 2013; Sivan et al., 2006). Additionally, oxidative stress participates in the intrinsic pathway of cellular apoptosis, and their roles have been confirmed in nucleus pulposus cells (NPCs) death and IDD induced by various risk factors(Dimozi, Mavrogonatou, Sklirou, & Kletsas, 2015; Feng et al., 2017; Suzuki et al., 2015). Meanwhile, various interventions targeting oxidative stress has shown beneficial effects on NPCs apoptosis and IDD(Hou, Lu, Chen, Yao, & Zhao, 2014; F. Wang, Cai, Shi, Wang, & Wu, 2016).

The endoplasmic reticulum (ER) is the largest organelle in cells and plays a vital role in many cellular process, such as polypeptide folding, calcium storage, lipid biosynthesis and secretion, as well as membrane protein maturation and transshipment(Schwarz & Blower, 2016). The endoplasmic reticulum is sensitive to and easily impaired by oxidative stress(Cubillos-Ruiz, Bettigole, & Glimcher, 2017; Dong et al., 2019; Görg et al., 2019). The production of misfolded protein exceeds the degradation under oxidative stress, which may cause the unfolded protein response (UPR), and further leads to the occurrence of endoplasmic reticulum stress (ER-stress)(Cao & Kaufman, 2014; Ron & Walter, 2007; Rutkowski & Kaufman, 2004).

Under physiological conditions, ER-transmembrane signaling molecules, such as protein kinase-like endoplasmic reticulum kinase (PERK), is inactive through binding to the ER-stress sensor, 78-kDa glucose-regulated protein (GRP78)(A. Lee, 2005). While in the condition of ER-stress, GRP78 is released from PERK. PERK is then autophosphorylated and activated. Activated PERK further phosphorylates the eukaryotic translation-initiation factor 2α (eIF2α) to attenuate protein synthesis(Cnop, Toivonen, Igoillo-Esteve, & Salpea, 2017; M. Wang, Wey, Zhang, Ye, & Lee, 2009). Then, when the stress is too severe to overcome, over-activated ER-stress promotes the expression of pro-apoptotic proteins, including C/EBP-homologous protein (CHOP)(M. Wang et al., 2009). Ultimately, under these conditions, cells undergo apoptosis. Studies have reported that ER-stress was significantly increased in NPCs during the IDD process(Lan, Shiyu-Hu, Shen, Yan, & Chen, 2021; Mizushima & Levine, 2020), while suppression ER-stress may lead to alleviated IDD(Liao et al., 2019; Xie et al., 2020; D. Xu et al., 2017).

ER-phagy was first discovered in yeast(Hamasaki, Noda, Baba, & Ohsumi, 2005), and described in mammalian cells in 2015(Khaminets et al., 2015). In the process of ER-phagy, part of the endoplasmic reticulum (ER) may be recognized by specific ER-phagy receptors, phagocytosed, and then transported to lysosomes to form autolysosomes, which are then degraded and removed(Nam & Jeon, 2019). Studies have reported that ER-phagy is involved in human neuropathy, degenerative diseases and various other diseases(Khaminets et al., 2015). Six ER-phagy receptors have been found in mammals so far, which including FAM134B, RTN3L, CCPG1, SEC62, TEX264, and ATL3(An et al., 2019; Chen et al., 2019; Chino, Hatta, Natsume, & Mizushima, 2019; Fumagalli et al., 2016; Smith et al., 2018; Wilkinson, 2019). Among which, FAM134B is the first discovered endoplasmic reticulum phagocytic receptor and demonstrated to potently promote ER-phagy in cells.

Nuclear factor-erythroid 2-related factor 2 (Nrf2) is known as a redox-sensitive basic leucine zipper transcription factor that regulates the expression of many antioxidants and phase II detoxifying enzymes. A number of studies have showed that activation of Nrf2 can inhibit the occurrence of ER-stress(Espinosa-Diez et al., 2015; Hernández-Gea et al., 2013; Liu et al., 2020; Mohamed et al., 2020) and alleviate a variety of diseases. At the meantime, as a transcription factor, it can significantly increase the level of autophagy(“Autophagy-Deficient Pancreatic Cancer Cells Depend on Macropinocytosis,” 2021; Dewanjee et al., 2021; Zhang, Feng, & Jiang, 2021). Therefore, we speculate that Nrf2 may be involved in the regulation of ER-phagy during oxidative stress.

In the current study, we reported for the first time that ER-phagy is decreased during oxidative stress in NPCs. Also, we observed that the expression of FAM134B was decreased, suggesting reduced ER-phagy during oxidative stress may due to decreased FAM134B expression. Furthermore, it was demonstrated that FAM134B mediated ER-phagy may protect NPCs against ER-stress and apoptosis. In addition, the nuclear transcription factor Nrf2 was found to promote the expression of FAM134B as well as ER-phagy, and protect NPCs against oxidative damage. *In vivo* study revealed that overexpression of FAM134B and Nrf2 could effectively attenuate the progression of IDD in rats. These results suggest Nrf2/FAM134B mediated ER-phagy may combat oxidative damage in cells; meanwhile, ER-phagy as well as Nrf2 could be potential therapeutic targets for IDD therapy.

## 2. Materials and Methods

### 2.1. Ethics statement

All surgical interventions, treatments and postoperative animal care procedures were strictly performed in accordance with the guidelines for Animal Care and Use outlined by the Committee of Wenzhou Medical University.

### 2.2. Confocal microscopy Scanning

Laser confocal experiments were acquired using a Zeiss LSM 800 confocal microscope equipped with a 63×1.4 numerical aperture oil objective. Airyscan microscopy was performed using a Zeiss LSM 800 confocal microscope, equipped with Plan-Apochromat 63×/1.4 numerical aperture oil objective and pixel size of 8.7 nm. Images were subjected to post-acquisition Airyscan processing. Image acquisition and processing were performed with Zen Blue software and co-localization analysis and image presentation was performed using Image J FIJI software or Photoshop (Adobe).

### 2.3. Primary rat NPCs isolation and culture

The lumbar segments (L1-L6) were extracted from Sprague-Dawley rats euthanized with an overdose of pentobarbital. Next, gel-like NP tissues were carefully collected under aseptic condition and transferred to culture dishes with 0.1% type II collagenase (Sigma-Aldrich, USA) treatment for 4 h at 37°C to digest the tissue completely. After a centrifugation at 1000 rpm for 5 min, the precipitate was resuspended and plated into high glucose Dulbecco modified Eagle medium (DMEM; Gibco, USA) with 25% fetal bovine serum (FBS; Gibco, USA) and 1% penicillin/streptomycin antibiotics (Gibco, USA). The isolated NPCs were kept in culture flasks in incubator maintained with 5% CO_2_ at 37°C. And the replacing of the complete medium was committed every other day. NPCs were exposed to tert-butyl hydroperoxide (TBHP; Sigma-Aldrich, USA) for in vitro induction of oxidative stress.

### 2.4. Lentivirus transfection

The lentiviral shRNA expression vector targeting LV-FAM134B, LV-shFAM134B, LV-Nrf2, LV-shNrf2 and RAMP-GFP-mCherry were produced by Genechem Technology (Shanghai, China). For transfection, the NPCs were seeded at 40e60% confluence and infected with lentivirus at a multiplicity of infection (MOI) of 80. After 12 h of transfection, the culture medium was changed every other day. When confluent, the transfected NPCs were passaged for further experiments. The transfection efficiency was evaluated by western blotting.

### 2.5. Western blotting

NPCs were lysed in ice-cold RIPA with 1 mM PMSF (phenylmethanesulfonyl fluoride, Beyotime). The protein concentration in the samples was measured using the BCA protein assay kit (Beyotime). The proteins were separated via sodium dodecyl sulfate–polyacrylamide gel electrophoresis and then transferred to a polyvinylidene difluoride membrane (Millipore, USA) followed by blocking with 5% nonfat milk. The bands were subsequently probed with primary antibodies. Finally, the intensity of the bands was quantified using Image Lab 3.0 software (Bio-Rad).

### 2.6. Lighting-Link Kit

Add 1 μL of Modifier reagent to each 10 μL of antibody to be labeled and mix gently. Remove cap from vial of DyLight Conjugation Mix and pipette the antibody sample (with added Modifier reagent) directly onto the lyophilized material. Resuspend gently by withdrawing and re-dispensing the liquid once or twice using a pipette. Replace cap on the vial and leave standing for 15 minutes in the dark at room temperature (20-25°C). After incubating for 15 minutes, add 1 μL of Quencher reagent for every 10 μL of antibody used and mix gently. The conjugate can be used after 5 minutes.

### 2.7. Immunofluorescence

NPCs were washed in PBS, fixed in 4% paraformaldehyde and permeated in 0.1% Triton X-100 for 15 min. The cells were then blocked with 5% bovine serum albumin for 1 h at 37 °C, rinsed with PBS and incubated with primary antibodies in a humid chamber overnight at 4 °C. The cells were washed and incubated with secondary antibodies for 1 h at room temperature and labeled with DAPI for 5 min. Images of each slide were obtained randomly with a fluorescence microscope (Olympus Inc., Tokyo, Japan). Images were prepared for presentation using Adobe Photoshop 6.0 (San Jose, CA, USA).

### 2.8. Quantitative RT-PCR

Total RNA was isolated using TRIzol (Invitrogen) according to the manufacturer’s instructions. cDNA was synthesized from 1μg of RNA via the One Step RT-PCR Kit (TaKaRa). Quantitative real time PCR was performed using the iQTM SYBR Green Supermix PCR kit with the iCycler apparatus system (Bio-Rad). GAPDH was used as the invariant housekeeping gene internal control.

### 2.9. Animal model

6 weeks old adult male Sprague Dawley rats were anesthetized by intraperitoneal injection with 2% (w/v) pentobarbital (40 mg/kg) and randomly divided into 5 groups (9 per group): Control, IDD+ LV-NC, IDD+ LV-FAM134B, IDD+ LV-Nrf2, IDD+ LV-Nrf2+ LV-shFAM134B. The whole layer of annulus fibrosus (AF) was vertically punctured through the skin of tail discs using the 21G needles after the localization of tail disc (Co7/8) by X-ray radiograph. To ensure that the needle puncture was not too deep, the length of the needle was decided according to the AF and the nucleus pulposus dimensions, which were measured in the preliminary experiment and found to be about 4 mm. All the needles were kept in the disc for 1 min. To eliminate the effect of injection volume on IDD, we injected LV-NC, LV-FAM134B, LV-shFAM134B or LV-Nrf2 into the center of the NP tissue using a 27G microliter syringe (Gaoge, China). After above surgical procedure, rats were monitored daily to observe their health condition.

### 2.10. Histological assessment

The rats were killed by an intraperitoneal overdose injection of 10% chloral hydrate and the tails were harvested. The human CEP tissues were collected during operation. The specimens were decalcified and fixed in formaldehyde, dehydrated and embedded in paraffin. The tissues were cut into 5-μm sections. Slides of each disc were stained with hematoxylin-eosin (HE staining) and Safranin-Orange (SO staining). Images were captured using a light microscope.

### 2.11. TUNEL staining assay

Treated NPCs were fixed with 4% paraformaldehyde for about 1 h. After being submerged with 3% H_2_O_2_ and 0.1% Triton X-100 for 10 min respectively, NPCs were washed by phosphate buffer saline (PBS) and stained by in situ Cell Death Detection Kit (Roche, USA) according to the manufacturer’s instructions. Additionally, 40, 6-diamidino-2-phenylindole (DAPI) was added to localize the nucleus. Finally, three random fields per slide were selected and captured by fluorescence microscope (Olympus Inc., Japan) to analyze the apoptosis of NPCs.

### 2.12. Transmission electron microscopy

After being fixed in 2.5% glutaraldehyde overnight, rat NPCs were fixed in 2% osmium tetroxide for 1 h and stained with 2% uranyl acetate for 1 h. Before embedding into araldite and cut into semithin sections, these samples were dehydrated in an ascending series of acetone. Then, semithin sections were stained with toluidine blue to locate cells before being observed with a transmission electron microscope (Hitachi, Tokyo, Japan).

### 2.13. Immunohistochemical analysis

The sections embedded in paraffin were deparaffinized and rehydrated while endogenous peroxidase was blocked by 3% hydrogen peroxide. The sections were incubated with 0.4% pepsin (Sangon Biotech, Shanghai, China) in 5 mM HCl at 37 °C for 20 min for antigen retrieval. The sections were incubated with 5% bovine serum albumin for 30 min at room temperature, followed by incubation with primary antibody overnight at 4 °C, and finally with HRP conjugated secondary antibody. The rate of positive cells in each section was quantitated by observers who were blinded to the experimental groups. Five mice of each group were used for quantitative analysis.

### 2.14. X-ray imaging acquisition

For X-ray imaging, it was performed for all rats at 0 week, 4 weeks, and 8 weeks after surgery by the X-ray irradiation system (Kubtec, USA). After imaging, the height of the intervertebral disc was measured using Image J software and the disc height index (DHI) was calculated to reflect the changes in intervertebral disc height as described in previous study.

### 2.15. Statistical analysis

Statistical analysis was performed using the Stata 20.0 software (IBM, Armonk, NY, USA). Data are expressed as the mean ± standard deviation (SD). Data were analyzed by one-way analysis of variance (ANOVA) followed by Tukey’s test for comparison between the two groups. Differences were considered significant when P < 0.05.

## 3. Results

### 3.1. The ER-phagy in nucleus pulposus cells under oxidative stress

The process of ER-phagy is briefly shown in figure 1A. LAMP1 is a membrane protein of lysosome, while Calnexin (CANX) is a chaperone protein that resides in the membrane of endoplasmic reticulum (ER). Thus, we used colocalization staining of fluorescently labeled LAMP1 and CANX to evaluate the level of ER-phagy, the immunofluorescence results were further quantified by the Pearson correlation coefficient. The results showed that the co-staining of LAMP1 and CANX was high in control group (physiological condition), while decreased during oxidative stress (TBHP treatment), and the effect was dose dependent (Fig. 1B&E).

**Figure 1.**
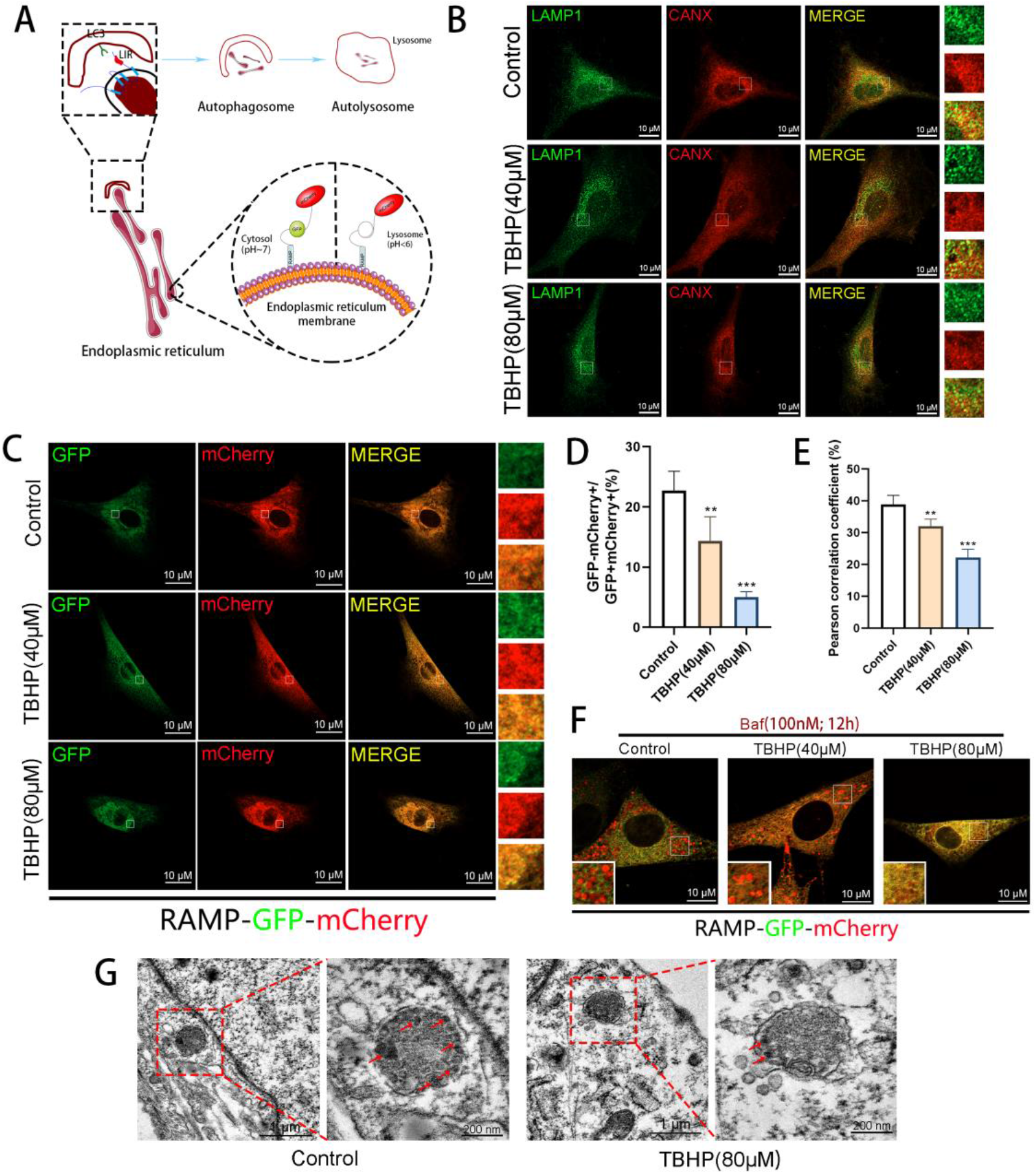
The ER-phagy in nucleus pulposus cells under oxidative stress (A) The process of ER-phagy and Schematic of the EATR assay. GFP is quenched as a result of low pH-induced protonation, causing a switch from GFP^+^/mCherry^+^ (yellow) to GFP^−^/mCherry^+^ (red) during ER-phagy. (B, E) Confocal scan analysis of LAMP1 (green) co-localization with CANX (red) in control or TBHP-exposed NPCs. Scale bar = 10 μm, n=5. (C, D) Measurement of the ER-phagy activity. NPCs transiently expressing RAMP-GFP-mCherry treated with TBHP. Scale bar = 10 μm, n=5. (F) The autolysosomal degradation of ER fragments labeled by RAMP-GFP-mCherry was blocked by Baf and the ER fragments labeled in lysosome can be observed easily under TBHP in various concentration. Scale bar = 10 μm, n=3. (G) TEM images of autophagic vesicles and ER-phagy in NPCs treated as indicated (Red arrow: parts of ER). Scale bar = 1μm or 200nm. All data represent mean ± S.D. \*\*\**P*< 0.001, \*\**P*< 0.01, relative to the control group.

To further assess the flux of ER-phagy during oxidative stress, RAMP-GFP-mCherry tandem reporter system was used in the study. This system was developed by Liang and colleagues(Liang, Lingeman, Ahmed, & Corn, 2018). In this system, RAMP4 is used as a marker of ER, and the fluorescence of GFP and mCherry were applied to indicate the localization of ER in cells. When the endoplasmic reticulum with a fluorescent tandem group is located in the cytoplasm, GFP and mCherry glow together, and the merge treatment shows yellow light (marked as GFP^+^ mCherry^+^); however, when ER-phagy occurs (ER is engulfed by lysosome), the acid environment in lysosome may induce the quenching of GFP, while retain mCherry (GFP^-^ mCherry^+^, observed by red). Thus, the ER-phagy flux could be evaluated according to the fluorescence of GFP and mCherry. For simplicity, we refer this system as ER autophagy tandem reporter (EATR; Fig.1A). Using this reporter system, it was found that the signal of GFP^-^ mCherry^+^ was high in control group, the treatment of TBHP resulted in decreased signal of GFP^-^ mCherry^+^ (Fig. 1C), and the quantification data showed that the ratio of GFP^-^ mCherry^+^/GFP^+^ mCherry^+^ was dose dependently decreased during TBHP treatment (Fig. 1C&D).

We then inhibit the activity of lysosome with the use of Bafilomycin A1 (Baf). Baf may inhibit the activity of various enzymes in the lysosome, while does not destroy the acidic environment inside the lysosome; thus, Baf may result in the accumulation of endoplasmic reticulum fragments and prevent further degradation, which makes ER-phagy easier to be observed. Subsequently, we simultaneously detected the expression of FAM134B and the intensity of ER-phagy through confocal microscope (Fig. 1F). Furthermore, we observed autophagosomes and selective autophagy by transmission electron microscopy (TEM). Compared to control group, TBHP-treated group showed less the autophagosomes contained parts of ER whorls (some rough ER parts with ribosomes adhesion) (Fig. 3G).

In summary, these results suggest that ER-phagy occurs in NPCs, and is decreased during oxidative stress.

### 3.2. FAM134B regulates the process of ER-phagy in nucleus pulposus cells

Next, we aimed to investigate how ER-phagy is decreased in NPCs during oxidative stress. FAM134B is a classic ER-phagy receptor (Fig. 2A) and a potent inducer of ER-phagy(Khaminets et al., 2015); thus, we evaluated the expression of FAM134B. *In vivo*, the expression of FAM134B was decreased in IDD, as shown by immunohistochemistry and immunofluorescence in rat nucleus pulposus tissues (Fig. 2B). Then, we evaluated the expression of FAM134B in NPCs by RT-PCR and western blot. The results showed the expression of FAM134B was dose dependently decreased during oxidative stress both at mRNA and protein level (Fig. 2C-E). Subsequently, we simultaneously detected the expression of FAM134B and the intensity of ER-phagy through a confocal microscope. The results showed that the FAM134B level and the co-localization degree of LAMP1 and CANX was decreased during oxidative stress (TBHP treatment), while was high in control group (physiological condition) (Fig. 2F). Finally, we use lentivirus to overexpress or knock down FAM134B, and detect the intensity of ER-phagy in nucleus pulposus cells by RAMP-GFP-mCherry tandem reporter system. The results showed that the knockdown of FAM134B resulted in the decrease of ER-phagy level in a physiological state, while the overexpression of FAM134B can up-regulate the level of ER-phagy. After TBHP treatment, the knockdown of FAM134B could lead to a further decline in ER-phagy level, while the overexpression of FAM134B can show a certain degree of recovery in ER-phagy level (Fig. 2G&H). TEM revealed that the less ER parts in autophagosomes in TBHP-treated NPCs; and when we treated NPCs with LV-FAM134B, the autophagosomes contained more ER parts (Fig. 2I). Finally, we used RAMP-GFP-mCherry and LV-FAM134B in NPCs. The process of ER-phagy was captured by confocal microscope (Fig. 2J).

**Figure 2.**
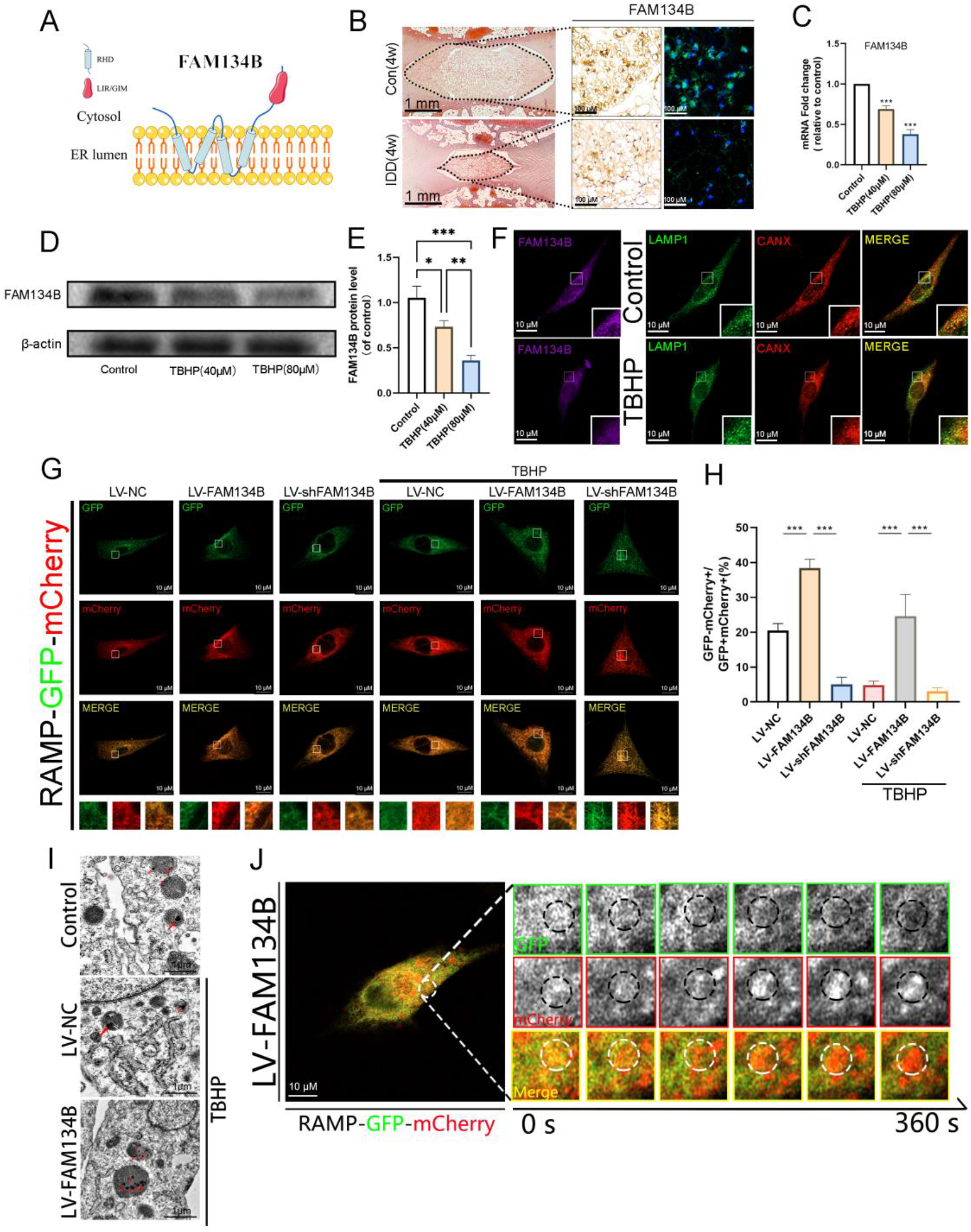
FAM134B regulates the process of ER-phagy in nucleus pulposus cells (A) A model of FAM134B. The classic ER-phagy receptor, bridging the gap between the ER and autophagic membranes. (B) The expression of FAM134B in the intervertebral disc tissue. Select the tissue immunohistochemistry and immunofluorescence results of the control group and IDD group to compare the expression of FAM134B. Scale bar = 1 mm or 100μm. (C) The gene expression of FAM134B was detected by RT-PCR in the NPCs, treated with or without the administration of TBHP, n=3. (D, E) The protein expression of FAM134B was detected by western blot in the NPCs, n=3. (F) The triple immunofluorescence staining of LAMP1 (green), CANX (red) and FAM134B (purple). Scale bar = 10 μm. (G&H) The comparison of the ER-phagy activity in different groups. Expressing RAMP-GPF-mCherry, NPCs were overexpressed or knocked down FAM134B via lentivirus and fixed to visualize GFP glowing under TBHP or not. Scale bar = 10 μm, n=5. (I) TEM images of autophagic vesicles and ER-phagy in NPCs treated as indicated (Red arrow: parts of ER). Scale bar = 1μm. (J) ER-phagy was captured by live imaging for 360 seconds. Scale bar = 10 μm. All data represent mean ± S.D. \*\*\**P*< 0.001, \*\**P*< 0.01, \**P*< 0.05, relative to the control group.

**Figure 3.**
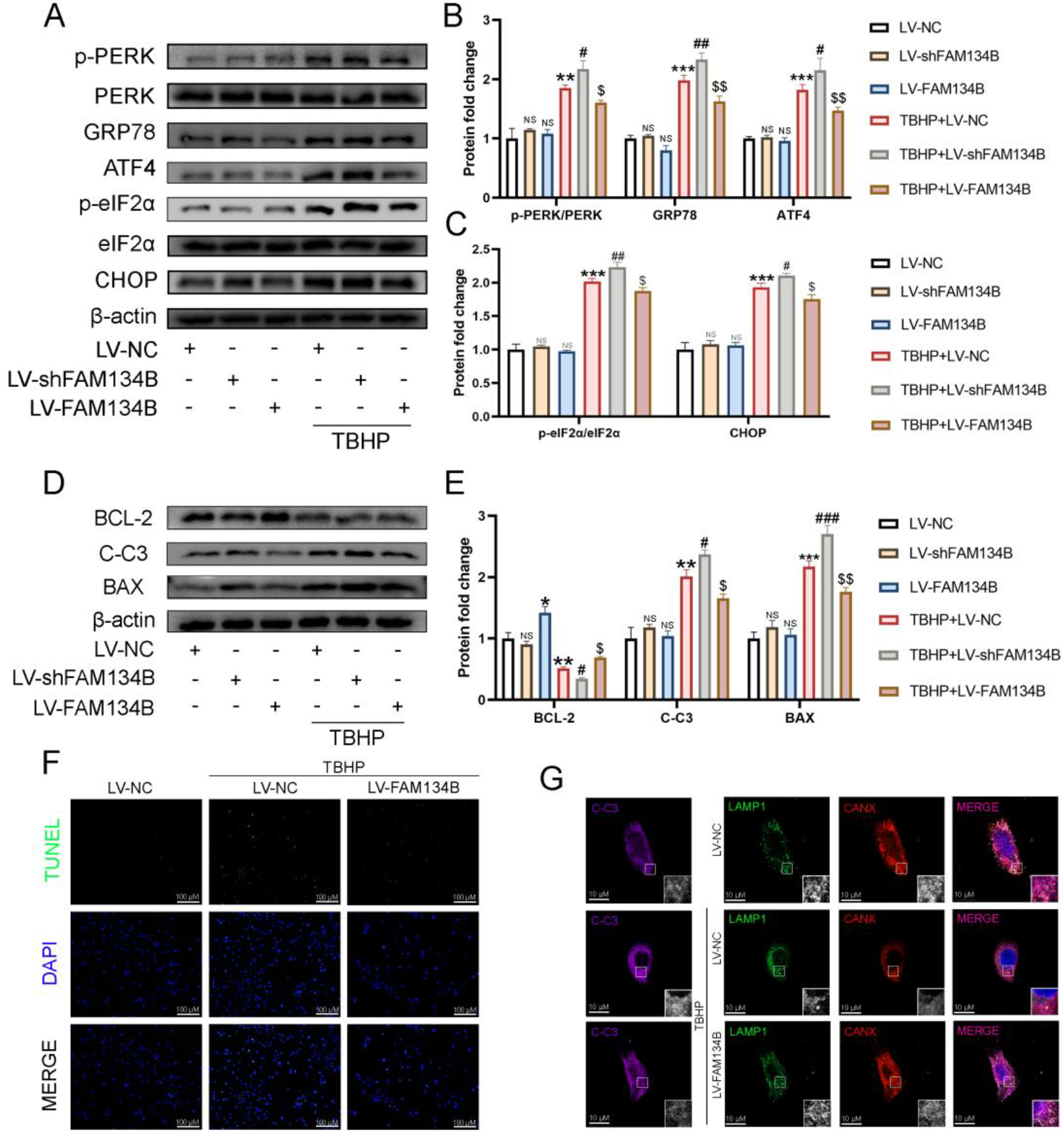
FAM134B suppresses ER-stress and apoptosis in nucleus pulposus cells (A, B, C) Measurement of the ER-stress level. The ER-stress related biomarker protein, such as p-PERK/PERK, GRP78, ATF4, p-eIF2α/eIF2α and CHOP were detected by western blot in the NPCs. (D, E) Measurement of the apoptosis in the NPCs. The protein expression of BCL-2, BAX and C-C3 detected by western blot. (F) Apoptotic NPCs were examined using TUNEL fluorescence immunocytochemistry (green). Nuclei were counterstained with DAPI (blue). Scale bar = 100 μm. (G) The triple immunofluorescence staining of LAMP1 (green)-CANX (red)-C-C3 (purple) in TBHP-exposed NPCs treated with LV-NC, LV-FAM134B. Nuclei were counterstained with DAPI (blue). Scale bar: 10 μm. All data represent mean ± S.D. All *in vitro* experiments were repeated three times independently. *\*\*\*P*< 0.001, \*\**P*< 0.01, \**P*< 0.05, relative to the control group. *###P*< 0.001, ##*P*< 0.01, #*P*< 0.05, relative to the TBHP-stimulated group. $$*P*< 0.01, $*P*< 0.05, relative to the TBHP+LV-shFAM134B group.

In summary, it is suggested that the expression of FAM134B is decreased during oxidative stress, while FAM134B can effectively regulate the level of ER-phagy in NPCs.

### 3.3. FAM134B suppresses ER-stress and apoptosis in nucleus pulposus cells

Next, we sought to evaluate the function of ER-phagy on ER-stress and apoptosis. Firstly, we detect the expression of PERK-ATF4-eIF2α-CHOP pathway by western blot, which can reflect the changes of ER-stress. We found that knocking down or overexpressing FAM134B under physiological conditions has no obvious effect on the PERK-ATF4-eIF2α-CHOP pathway proteins. However, when NPCs are oxidatively stressed (TBHP treatment), the knockdown of FAM134B may increase the intensity of ER-stress, and the overexpression of FAM134B can effectively alleviate ER-stress (Fig. 3A-C).

Subsequently, we detected the expression of apoptosis-related proteins such as Bcl-2, Cleaved-Caspase 3 (C-C3), and BAX. Similar to the expression of PERK-ATF4-eIF2α-CHOP pathway protein mentioned above, the knockdown and overexpression of FAM134B under physiological conditions cannot cause the activation of the apoptosis pathway. However, after TBHP treatment, the knockdown of FAM134B increased the apoptosis of NPCs, and the overexpression of FAM134B reduced the apoptosis of NPCs stimulated by oxidative stress (Fig. 3D&E). Subsequently, we detect the effect of overexpression of FAM134B on apoptosis through TUNEL staining, and we can get the same experimental conclusion as the previous one (Fig. 3F). In addition, LAMP1 (488nm)-CANX (594nm)-C-C3 (647nm) triple fluorescence staining was used by lighting link kit, and it was found that after TBHP stimulation treatment, the degree of colocalization of LAMP1-CANX decreased, and the fluorescence intensity of C-C3 increased, which indicates increased apoptosis of NPCs; after LV-FAM134B overexpression treatment, the degree of co-localization of LAMP1-CANX increased, and the fluorescence intensity of C-C3 decreased, which shows that the apoptotic damage of nucleus pulposus cells is rescued (Fig. 3G).

In summary, these results show that FAM134B medicate ER-phagy may suppresses ER-stress and apoptosis in NPCs during oxidative stress.

### 3.4. FAM134B exerts its effect in nucleus pulposus cells under oxidative stress through ER-phagy

In the next step, we aimed to prove that FAM134B exerts its effects by regulating ER-phagy. We use Baf as an inhibitor of ER-phagy to observe whether the therapeutic effect of FAM134B is through ER-phagy. According to the results of confocal microscopy, the ER-phagy level of NPCs decreased under the stimulation of TBHP, and the ER-phagy intensity increased significantly after overexpression of FAM134B protein. When Baf was used to treat NPCs which were treated with LV-FAM134B, obvious lysosome accumulation was observed in the cells, suggesting that the ER-phagy process was blocked (Fig. 4A). After that, we used western blot to verify that the expression of apoptosis-related protein and ER-stress-related proteins. After Baf treatment, the therapeutic effect of FAM134B overexpression was suppressed (Fig.4B-G). Finally, we used CHOP (488nm) and BCL-2 (594nm) immunofluorescence double staining and TUNEL staining to verify and draw similar conclusions (Fig.4H&I).

**Figure 4.**
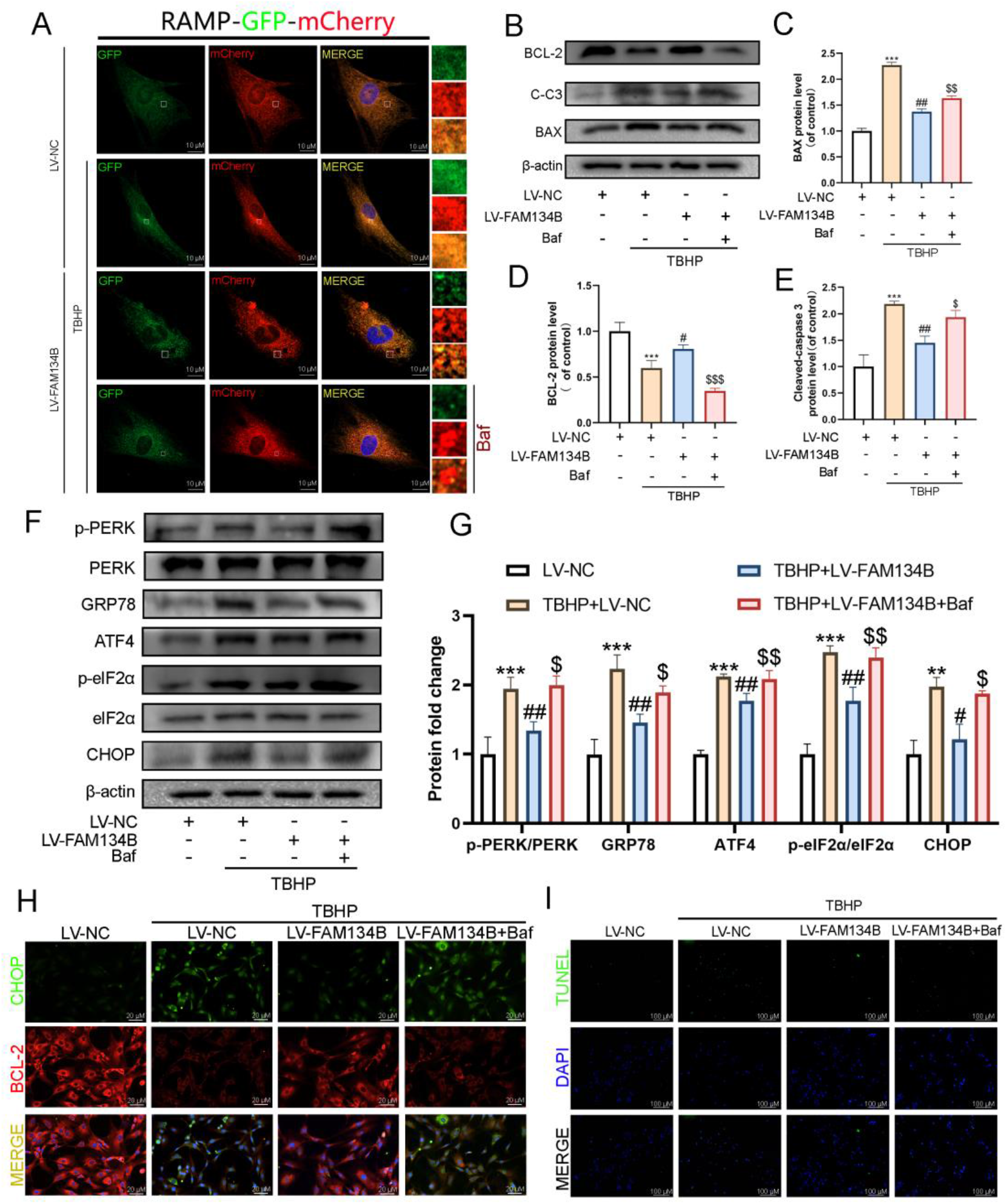
FAM134B exerts its effect in nucleus pulposus cells under oxidative stress through ER-phagy (A) Measurement of the ER-phagy activity. NPCs transiently expressing RAMP-GFP-mCherry were treated with Baf as inhibitor. Scale bar = 10 μm. (B-E) Measurement of the apoptosis in the NPCs. The protein expression of BCL-2, BAX and C-C3 detected by western blot. (F, G) Western blot analysis of ER-stress level. The protein expression of p-PERK/PERK, GRP78, ATF4, p-eIF2α/eIF2α and CHOP were detected by western blot in the NPCs. (H) Immunofluorescence staining of CHOP (green) and BCL-2 (red). Nuclei were counterstained with DAPI (blue). Scale bar: 20 μm. (I) Apoptotic NPCs were examined using TUNEL fluorescence immunocytochemistry (green). Nuclei were counterstained with DAPI (blue). Scale bar: 100 μm, n=3. All data represent mean ± S.D. All *in vitro* experiments were repeated three times independently. *\*\*\*P*< 0.001, \*\**P*< 0.01, \**P*< 0.05, relative to the control group. ##*P*< 0.01, #*P*< 0.05, relative to the TBHP-stimulated group. $$*P*< 0.01, $*P*< 0.05, relative to the TBHP+LV-FAM134B group.

Together, these results suggest that FAM134B exerts its effects by regulating ER-phagy.

### 3.5. Nrf2 promotes the expression of FAM134B and ER-phagy

The results above show that ER-phagy may suppress oxidative damage in NPCs; however, there is no specific agonist for ER-phagy so far. Thus, we aimed to explore the upstream regulator of ER-phagy from the view of FAM134B. Nuclear transcription factor Nrf2 is closely related to the level of oxidative stress. Nrf2 has been proven to effectively inhibit oxidative stress in cells. So, we asked whether Nrf2 may regulate ER-Phagy.

Firstly, we use LV-Nrf2 and LV-sh-Nrf2 to overexpress or knock down the mRNA expression of Nrf2. According to the RT-PCR results, we found that when LV-Nrf2 is used to overexpress the level of Nrf2 mRNA level in NPCs, the mRNA level of FAM134B will also be further upregulated. When Nrf2 was knocked down by LV-shNrf2, FAM134B mRNA will reduce (Fig. 5A). Then, we use western blot to verify the effect of LV-shNrf2 and LV-Nrf2 shows that LV-shNrf2 can knock down the protein levels of total and nuclear Nrf2, and after using LV-Nrf2, the protein levels of total and nuclear Nrf2 are significantly increased. It is proved that lentivirus can effectively regulate the protein expression of Nrf2 (Fig. 5B&C). Meanwhile, we used lentivirus to adjust the expression level of Nrf2 to observe the changes of FAM134B by western blot. Results show that Nrf2 knockdown can reduce FAM134B expression; when Nrf2 over-expression, FAM134B can be up-regulated (Fig. 5B&C). Therefore, we have reason to believe that Nrf2 has the potential to regulate FAM134B. Finally, we used a confocal microscope to observe the changes in the phagocytic level of NPCs after the lentivirus overexpressed Nrf2 or knocked down Nrf2 under physiological condition or TBHP treatment (Fig. 5D&E).

**Figure 5.**
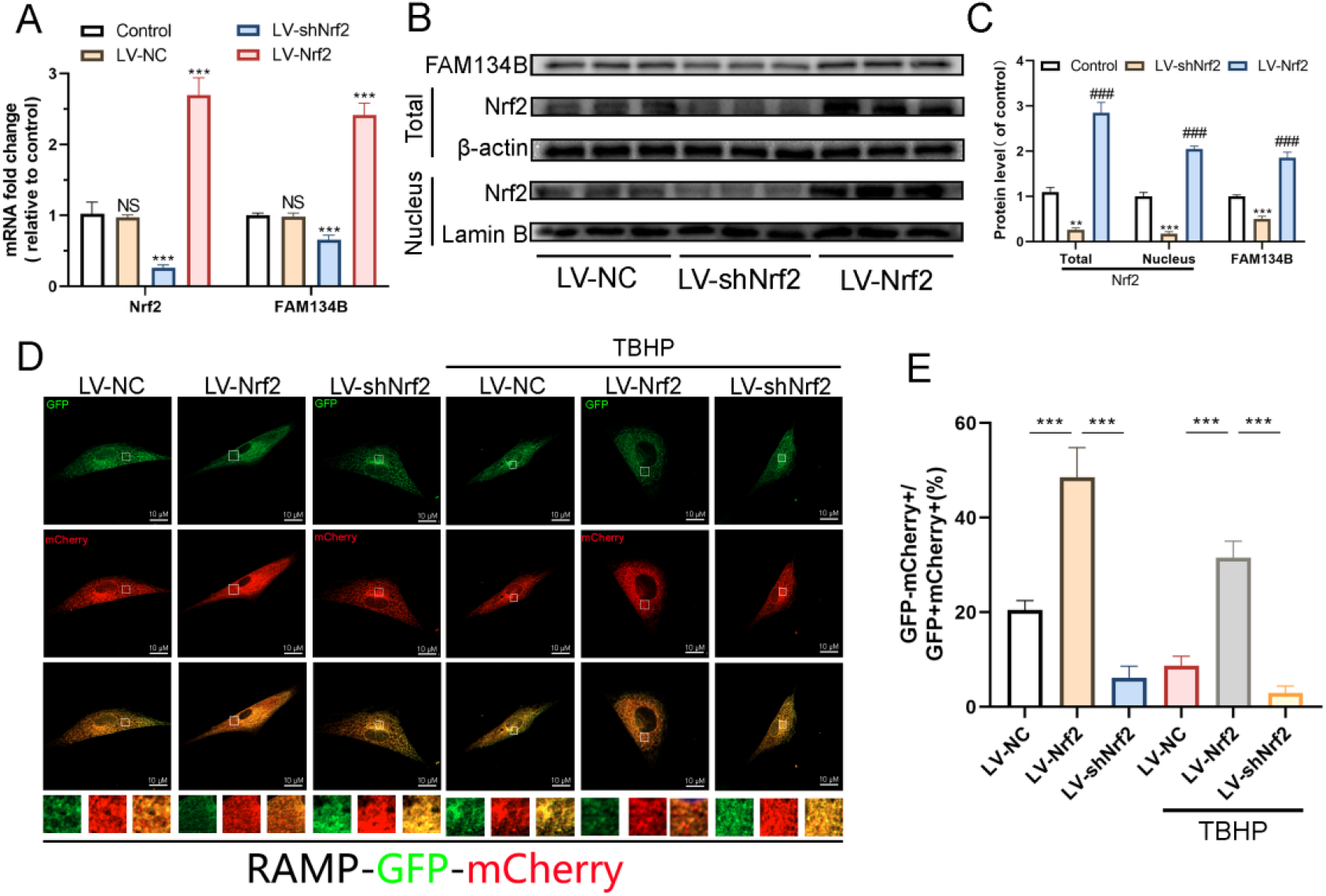
Nrf2 promotes the expression of FAM134B and ER-phagy (A) The gene expression of FAM134B and Nrf2 were detected by RT-PCR in the NPCs, treated with the LV-Nrf2 or LV-shNrf2, n=3. (B, C) Evaluation of the level of Nrf2 in the nucleus and in the whole cells by western blot, treated with or without the LV-NC and LV-shNrf2. And the expression of FAM134B treated with or without the LV-NC and LV-shNrf2 by western blot, n=3. (D&E) Confocal microscopy observed the changes in ER-phagy intensity in NPCs treated with LV-shNrf2 and LV-Nrf2 under TBHP treated or not. Scale bar: 10 μm, n=5. All data represent mean ± S.D. *\*\*\*P*< 0.001, relative to the control group. *###P*< 0.001, relative to the LV-Nrf2-treated group.

Our results show that Nrf2 can effectively regulate the expression of FAM134B, thereby affecting the activity of ER-phagy.

### 3.6. Nrf2 suppresses ER-stress and apoptosis through FAM134B mediated ER-phagy

Next, we sought to prove that whether Nrf2 can exert its effect on ER-stress and apoptosis through ER-phagy regulation. Firstly, we use western blot to detect the level of FAM134B, ER-stress and apoptosis-related proteins in NPCs after overexpression or knockdown of Nrf2 with lentivirus. The results showed that when the NPCs were treated by TBHP, the expression of ER-stress-related proteins in the NPCs increased, and at the same time, apoptosis further increased. With the knockdown of Nrf2, this phenomenon becomes more obvious. Oppositely, when we overexpressed Nrf2, the treatment effect of TBHP was effectively inhibited (Fig. 6A-D). Subsequently, we used CHOP (488nm)-BCL-2 (594nm) immunofluorescence double staining to detect the level of ER-stress and apoptosis. The results showed that knockdown of Nrf2 resulted in an increase in CHOP expression and a decrease in BCL-2 expression, while overexpression of Nrf2 inhibited this phenomenon (Fig.6E).

**Figure 6.**
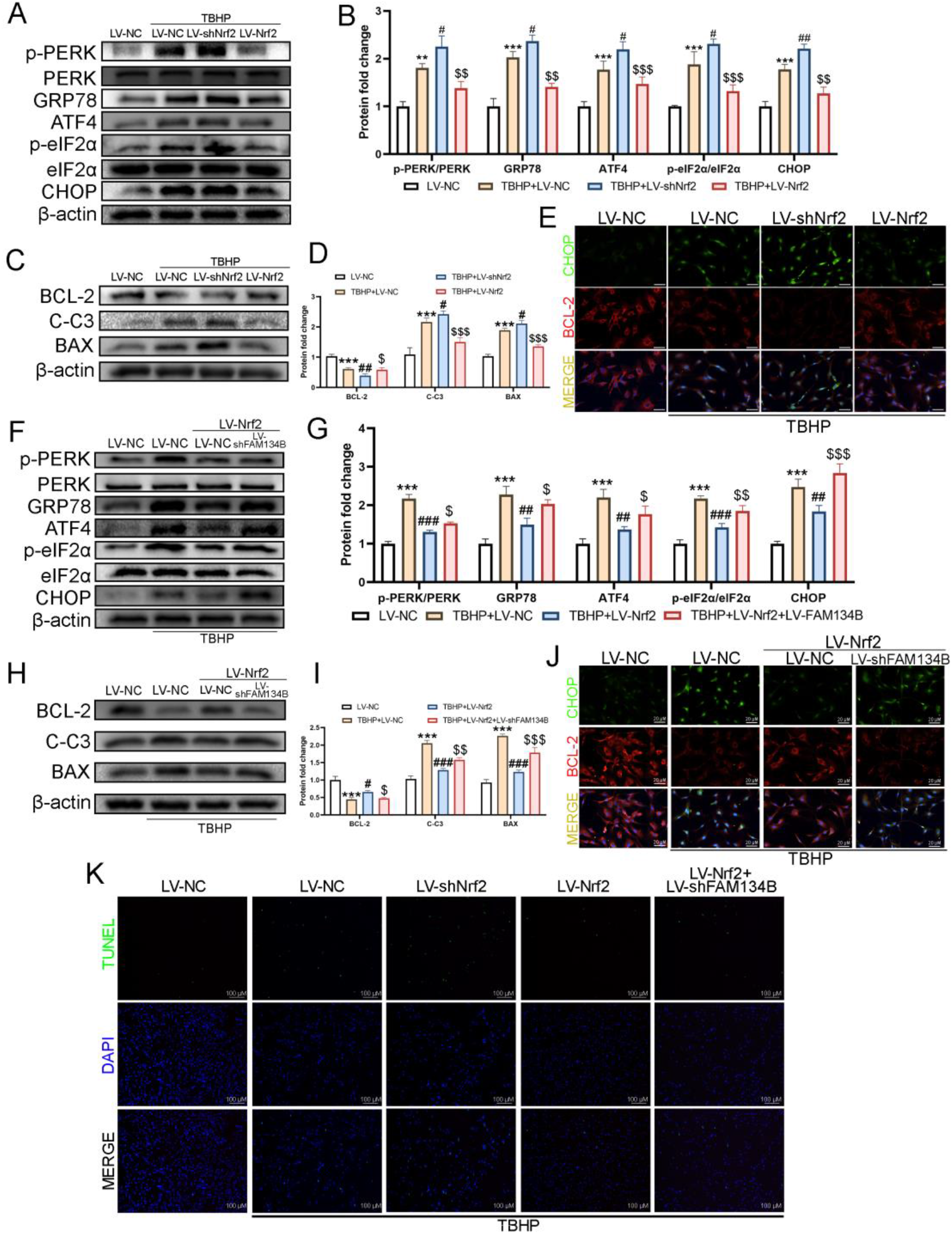
Nrf2 suppresses ER-stress and apoptosis through FAM134B mediated ER-phagy (A, B) Western blot analysis of the ER-stress level. The protein expression of p-PERK/PERK, GRP78, ATF4, p-eIF2α/eIF2α and CHOP were detected by western blot in the NPCs. (C, D) Measurement of the apoptosis in the NPCs. The protein expression of BCL-2, BAX and C-C3 detected by western blot. (E) Immunofluorescence staining of CHOP (green) and BCL-2 (red) in different groups. Nuclei were counterstained with DAPI (blue). Scale bar: 20 μm. (F, G) Measurement of the ER-stress level in the NPCs. The protein expression of p-PERK/PERK, GRP78, ATF4, p-eIF2α/eIF2α and CHOP were detected by western blot. (H, I) Western blot analysis of apoptosis in the NPCs. The protein expression of BCL-2, BAX and C-C3 detected by western blot. (J) Immunofluorescence staining of CHOP (green) and BCL-2 (red) in different groups. Nuclei were counterstained with DAPI (blue). Scale bar: 20 μm. (K) Apoptotic NPCs were examined using TUNEL fluorescence immunocytochemistry (green). Nuclei were counterstained with DAPI (blue). Scale bar: 100 μm. All data represent mean ± S.D. All *in vitro* experiments were repeated three times independently. *\*\*\*P*< 0.001, \*\**P*< 0.01, relative to the control group. *###P*< 0.001, ##*P*< 0.01, #*P*< 0.05, relative to the TBHP-stimulated group. $$*P*< 0.01, $*P*< 0.05, relative to the TBHP+LV-shNrf2 or TBHP+ LV-shNrf2 group.

After that, we verified by western blot that the therapeutic effect induced by Nrf2 can be reversed after LV-shFAM134B knockdown. As expected, after knocking down of FAM134B, the ER-stress and apoptosis biomarkers increased significantly, comparing with LV-Nrf2 treatment group (Fig.6F-I). Immediately afterwards, the immunofluorescence double staining results of CHOP (488nm)-BCL-2 (594nm) also confirmed the same phenomenon (Fig. 6J). TUNEL results also confirmed the same point of view (Fig.6K).

In summary, these results indicate that Nrf2 may suppress ER-stress and apoptosis through FAM134B mediated ER-phagy.

### 3.7. Overexpression of FAM134B and Nrf2 ameliorates intervertebral disc degeneration in rats in vivo

Based on the results of the *in vitro* experiments, we further studied the role of ER-phagy and Nrf2 *in vivo*. To confirm the therapeutic effect of FAM134B andNrf2 in IDD *in vivo*, we injected lentivirus into NP tissue to regulate the expression of FAM134B and Nrf2, and assess its effects on IDD progress in rats.

The intervertebral discs were evaluated at 0, 4, and 8 weeks after IDD surgery. We used the disc height index (DHI) to evaluate the degree of IDD based on images obtained by X-ray, respectively (Fig.7A, B). The height of the intervertebral disc displayed by X-ray was normal, and there was no significant difference among groups at 0 week after IDD surgery. However, at 4 and 8 weeks after IDD surgery, the IDD group had lower disc height compared with the control group. Interestingly, FAM134B and Nrf2 overexpression delayed the loss of intervertebral disc height. But when FAM134B is knocked down, it can inhibit the therapeutic effect brought by overexpression of Nrf2.

**Figure 7.**
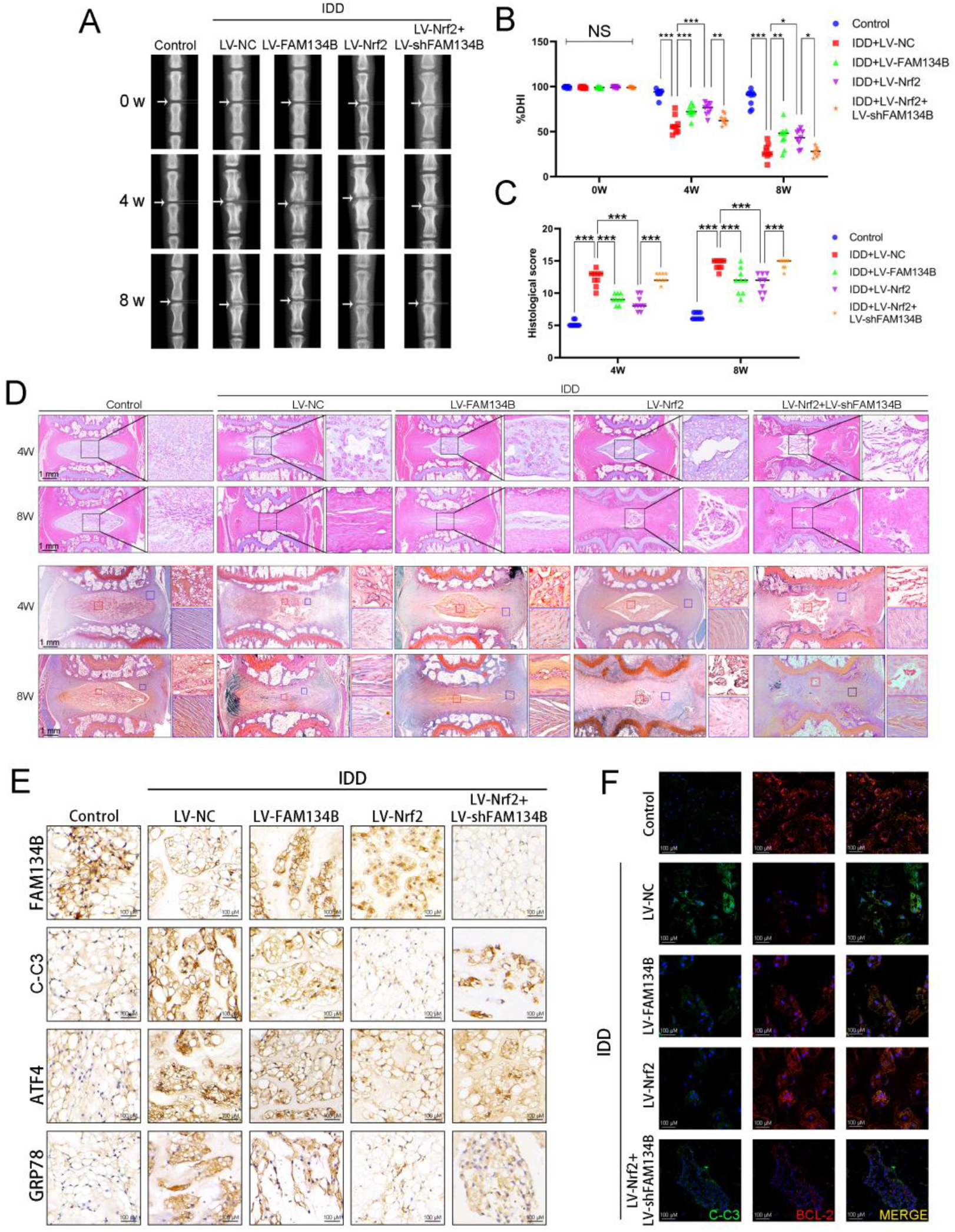
Overexpression of FAM134B and Nrf2 ameliorates intervertebral disc degeneration in rats *in vivo* (A, B) X-rays of the intervertebral disc in rats from the different experimental groups (Sham group, IDD+LV-NC group, IDD+LV-FAM134B group, IDD+LV-Nrf2 group and IDD+LV-Nrf2+LV-shFAM134B group), and loss of DHI was found (white arrows). (C, D) Representative HE and SO staining of the intervertebral disc (scale bar: 1 mm). (E) Immunohistochemical staining assay for FAM134B, C-C3, ATF4 and GRP78 in the NP tissue. Scale bar: 100μm. (F) Immunofluorescence staining of C-C3 (green) and BCL-2 (red) in different groups. Nuclei were counterstained with DAPI (blue). Scale bar: 100 μm. All data represent mean ± S.D, *n*=9. \*\*\**P*< 0.001, \*\**P*< 0.01, \**P*< 0.05.

From histomorphology assessed by Hematoxylin-Eosin (HE) staining and Safranine O-Fast Green (SO) staining, it could be found that with the process of IDD, the NPCs of the IDD group were replaced by fibrochondrocytes, the NP volume shrunk significantly until disappeared, and the annulus fibrosus (AF) structure was severely impaired. In addition, SO staining suggested that the glycosaminoglycan and proteoglycan content of NP were also decreased over time. However, FAM134B and Nrf2 overexpression lentivirus treatment significantly delayed the development of IDD (Fig.7C&D).

To investigate the treatment of FAM134B and Nrf2 *in vivo*, we detect the expression of FAM134B, C-C3, GRP78 and ATF4 by the immunohistochemical staining. It was proved that overexpression of FAM134B and Nrf2 can attenuate the expression of ER-stress and apoptosis-related proteins *in vivo* (Fig.7E). At the same time, it has been confirmed again that knockdown of FAM134B can weaken the therapeutic effect of Nrf2. The immunofluorescence double staining results of C-C3 (488nm)-BCL-2 (594nm) also confirmed the same phenomenon (Fig.7F).

These results demonstrate that overexpression of FAM134B and Nrf2 could ameliorate IDD in rats *in vivo*.

## 4. Discussion

In the current study, we reported for the first time that the ER-phagy level is reduced during oxidative stress. Through manipulating ER-phagy by FAM134B, we found that ER-phagy may suppress ER-stress and apoptosis in NPCs. At the meantime, it was also found that Nrf2 may regulate the expression of FAM134B, thus promote ER-phagy in NPCs.

Through co-localization of LAMP1 and CANX, as well as RAMP-GFP-mCherry system, we found the level of ER-phagy is significantly reduced under oxidative stress (TBHP stimulation) (Fig. 1). Liang, J *et al*. investigated ER-phagy in nutrient deprivation condition, they cultured cell in Earl’s buffered saline solution (EBSS) and found that it could potently induce ER-phagy(Liang et al., 2020). Different from previous studies, we observed oxidative stress may also regulate ER-phagy, though it may not promote, but suppress ER-phagy in NPCs.

Next, we explored the mechanism of the reduced ER-phagy in NPCs during oxidative stress. FAM134B (also known as JK-1, RETREG1) is the first endoplasmic reticulum phagocytic receptor identified. In 2001, FAM134B was first identified as an oncogene in esophageal squamous cell carcinoma (ESCC)(Tang, Lam, Law, Wong, & Srivastava, 2001). In the more than 20 years since its discovery, the biological functions of FAM134B have gradually been revealed. In recent years, FAM134B dysfunction has been reported to be involved in many diseases, including neuropathy, viral infection(Chiramel, Dougherty, Nair, Robertson, & Best, 2016; Lennemann & Coyne, 2017), vascular disease(Kong, Kim, & Lee, 2011; Li, Chen, Li, Yao, & Liu, 2021), inflammation(Yao, Ren, Xia, & Yao, 2021) and cancers(K. Lee et al., 2020). FAM134B has a LC3 interaction domain (LIR); thus, it may potently induce ER-phagy and participate in many ER-phagy-related processes, such as quality control of procollagens and endoplasmic reticulum-lysosomal associated degradation (ERLAD).

We found that both mRNA and protein level of FAM134B was significantly down-regulated during oxidative stress, and this phenomenon was also confirmed in degenerative nucleus pulposus tissues in rats (Fig. 2). These results suggested that the reduced level of ER-phagy in NPCs may due to the down-regulated expression of FAM134B during oxidative stress. Besides FAM134B, lots of proteins have been reported to regulate ER-phagy in cells, such as TEX264(Chino et al., 2019), SEC62(Fumagalli et al., 2016), CCPG1(Smith et al., 2018),RTN3L(Zou et al., 2018), and ATL3(Chen et al., 2019); thus, it was possible that other proteins may also be involved in the reduced ER-phagy under oxidative stress, which of cause need further validation.

The regulation of oxidative stress on ER-phagy is the main finding of our study. Besides our results, we think there may be other mechanisms that oxidative stress regulates ER-phagy. Mitochondria are sensitive to oxidative stress, and mitochondrial oxidative phosphorylation (OXPHOS) is reported to be impaired under oxidative stress(Bai et al., 2011; López-Erauskin et al., 2012; Satapati et al., 2016; Schulz et al., 2007; Zorov, Juhaszova, & Sollott, 2014). Meanwhile, disruption of mitochondrial OXPHOS system may inhibit ER-phagy (Liang et al., 2020). We hypothesized that TBHP induced oxidative stress may lead to mitochondrial OXPHOS damage, thereby affect the activity of ER-phagy in NPCs.

Meanwhile, we focus on the function of ER-phagy during oxidative stress, from the view of the relationship between ER-phagy and ER-stress. Our study showed that upregulating ER-Phagy through over-expression of FAM134B can effectively relieve ER-stress as well as its downstream effect (apoptosis) under oxidative stress (Fig. 3&4). At the same time, defective ER-phagy can promote the occurrence of ER-stress (Fig.3&4).

ER is a membrane-bound organelle that is specialized for the folding and post-translational maturation of almost all membrane proteins and most secreted proteins(Cao & Kaufman, 2014). During oxidative stress, the unfolded and misfolded protein may be increased, which may further lead to ER-stress(Hetz, Martinon, Rodriguez, & Glimcher, 2011; Kawaguchi & Ng, 2011). As unfolded proteins accumulate, the ER becomes swollen and eventually fragmented(Bernales, Soto, & McCullagh, 2012). ER-phagy is a process that the ER fragments are engulfed by autophagosomes and degraded by lysosomes(Dikic, 2017; Mochida et al., 2015). Thus, the suppressive effect of ER-phagy on ER-stress may due to the clearance of ER fragments, which are the inducer of ER-stress. The effect of ER-phagy on ER-stress has also been proved by other studies(Chen et al., 2019; Delorme-Axford, Popelka, & Klionsky, 2019; Forrester et al., 2019).

Furthermore, we also found that the transcription factor Nrf2 may regulate the expression of FAM134B. When we knocked down Nrf2 with lentivirus, the expression level of FAM134B and ER-phagy was significantly reduced; and when we over-expressed Nrf2, it was able to promote the expression of FAM134B and ER-phagy, both in physiological and oxidative stress conditions (Fig.5&6). The expression of FAM134B has been reported to be regulated by transcription factor MiT/TFE(Cinque et al., 2020); our study showed for the first time that Nrf2, also as a transcription factor, may regulate the expression of FAM134B as well.

However, it is interesting that the expression of Nrf2 was reported to be increased under oxidative stress in the cells(J. Li et al., 2021; X. Xu et al., 2021), although not in NPCs, which seems to be contrary to our findings. According to the existing results, the expression of Nrf2 is controlled both by Nrf2 and MiT/TFE factors(Cinque et al., 2020). In our previous study, it was found that the nuclear expression of TFEB was reduced in NPCs under oxidative stress(Zheng et al., 2019). We propose that TFEB may have greater potential than Nrf2 to regulate the expression of Nrf2; thus, the reduced expression of FAM134B may mainly due to the decreased activity of TFEB.

In *in vivo* study, it was shown that both overexpression of FAM134B and Nrf2 may postpone the progression of IDD in rats, suggesting both ER-phagy and Nrf2 may serve as therapeutic targets for IDD. There is no specific ER-phagy or FAM134B agonist so far; however, plenty of compounds as well as FDA approved drugs have been reported to activate Nrf2(Ashrafian et al., 2012; Bae et al., 2013; “Bayer’s Vividion Buy Targets Undruggable Proteins,” 2021; Mills et al., 2018). Therefore, we suppose that Nrf2 agonists may have great potential to treat ER-phagy deficient conditions.

## Acknowledgments

This study was funded by Zhejiang Provincial Natural Science Foundation of China (LGF21H060011, LGF20H060013, LGF21H060010, LQ19H060004), National Natural Science Foundation of China (82172494, 81871806, 81972094, 81902243), Wenzhou Science and Technology Bureau Foundation (ZY2019014, Y2020059, Y20180031), Lin He’s New Medicine and Clinical Translation Academician Workstation Research Fund (18331213).

## Authors’ contributions

Zhen Lin, Xiangyang Wang and Xiaolei Zhang designed the project. Cheng Teng, Zhao Zhang, Xinlei Lu, Yaosen Wu and Long Wu performed the experiments and analyzed the results. Yifei Zhou, Liaojun Sun, Naifeng Tian and Zhongke Lin revised the manuscript. Zhen Lin, Zongyou Pan, Xiangyang Wang, and Libin Ni drafted the manuscript and Chenglong Xie and Xinlei Lu edited languages as well as grammatical concerns of the full manuscript. Finally, all authors successfully revised the final version of the manuscript.

